# Pairwise sequence similarity mapping with PaSiMap: reclassification of immunoglobulin domains from titin as case study

**DOI:** 10.1101/2022.05.13.491469

**Authors:** Kathy Su, Olga Mayans, Kay Diederichs, Jennifer R. Fleming

**Affiliations:** Department of Biology, Universität Konstanz, Konstanz, Baden Württemberg, 78456, Germany

## Abstract

Sequence comparison is critical for the functional assignment of newly identified protein genes. As uncharacterised protein sequences accumulate, there is an increasing need for sensitive tools for their classification. Here, we present a novel multidimensional scaling pipeline, PaSiMap, which creates a map of pairwise sequence similarities. Uniquely, PaSiMap distinguishes between unique and shared features, allowing for a distinct view of protein-sequence relationships. We demonstrate PaSiMap’s efficiency in detecting sequence groups and outliers using titin’s 169 immunoglobulin (Ig) domains. We show that Ig domain similarity is hierarchical, being firstly determined by chain location, then by the loop features of the Ig fold and, finally, by super-repeat position. The existence of a previously unidentified domain repeat in the distal, constitutive I-band is revealed. Prototypic Igs, plus notable outliers, are identified and thereby domain classification improved. This re-classification can now guide future molecular research. In summary, we demonstrate that PaSiMap is a sensitive tool for the classification of protein sequences, which adds a new perspective in the understanding of inter-protein relationships. PaSiMap is applicable to any biological system defined by a linear sequence, including nucleotides.

## INTRODUCTION

Genomic databases continue to grow at an accelerated pace generating a pressing need to effectively classify and analyze protein sequences. At the time of writing, there are nearly 220 million sequences in the UniProtKB database, of these only 565 thousand are contained in UniProtKB/Swiss-Prot, the section archiving manually annotated records. This leaves 99.7% of known sequences annotated using automated pipelines. Currently, the most reliable and effective way for automated pipelines to computationally interpret the sequence of an uncharacterized protein is by establishing its homology to proteins of known structure and function. However, establishing such relationships accurately can be challenging, particularly as large protein families often consist of multiple subclasses that can trouble the assignment of distant homologues with non-canonical features. Grouping proteins by sequence similarity is commonly the first step in inferring homology and, thereby, biological function. However, the accuracy of such classification closely depends on the quality of the sequence grouping.

Traditionally, phylogenetic trees were used to derive such information, but their reliance on underlying evolutionary models, such as linear relationships between sequences, and difficulties in visualizing large datasets have led to them becoming less favored. In recent years, network-based and multidimensional scaling of protein sequence inter-relationships have been explored for identifying sequence groupings, in conjunction with or as a replacement to phylogenetic trees (1–5). These methodologies can be split into two main approaches: those that use multiple sequence alignments (MSAs) and those independent of this requirement. Generally, it is considered advantageous to avoid MSAs as they can introduce bias, become cumbersome for large datasets, and often require user curation to produce meaningful results (1). Multidimensional scaling procedures fill a need not addressed by conventional phylogenetic tree construction, namely speed and reproducibility of calculations in large datasets, the display of all and not just optimal scoring connections and the ease of visualizing relationships - which are often obscured by complex tree structures (1). To advance the sensitivity and offer a novel perspective on protein sequence classification, we have developed a new multidimensional scaling pipeline which identifies sequence groups that share a distinguishing common feature. The approach reveals canonical prototypic group members and is particularly sensitive at identifying group outliers. The pipeline not only maps individual sequences as data points according to their pairwise similarities but also organizes the sequences based on shared and unique features within each group, thereby communicating which group sequences belong to as well as how far each sequence deviates from the core features of that group. At its core, the algorithm utilizes a multidimensional scaling method, *cc_analysis* (6), which was originally developed for the processing of experimental diffraction data in X-ray crystallography, and applies it to the problem of protein sequence grouping, creating a sequence analysis pipeline: <u>Pa</u>irwise <u>Si</u>milarity <u>Map</u> (PaSiMap). This pipeline aligns all possible pairs of sequences. For each resulting pair alignment, a similarity value is calculated. This produces a matrix of pairwise similarities which is then fed into *cc_analysis*, which represents these similarities and differences as coordinates in low-dimensional (e.g. 2D or 3D) space. The application scope of PaSiMap is not limited to partitioning sequences into discrete groups. Instead, it provides information on a continuous level, such as whether a pair of sequences (or groups) are more similar in comparison to another sequence (or group). PaSiMap is deterministic since it does not depend on a random number generator therefore it gives the same output for any given input; it is also robust as the result is not significantly influenced by the addition or removal of individual sequences. PaSiMap offers an alternative data visualization and grouping approach that can contribute to protein classification and the development of new biological hypotheses.

To demonstrate the sensitivity of PaSiMap for the identification of protein groups and the detection of outliers, we have selected a set of closely related sequences corresponding to the immunoglobulin domains of the protein titin from the muscle sarcomere. Titin is a molecular giant that spans half sarcomeres, from the Z-disk to the M-line (Fig 1A), acting to preserve passive force and coordinating sarcomere mechano-signaling (7, 8). Despite its diverse and intricate functionalities in muscle, the molecular composition of titin is simple and repetitive, consisting of over 300 homologous Immunoglobulin (Ig) and Fibronectin (Fn) domains (each *ca* 100 amino acids in length) linked in tandems and in occasions organized into super-repeats (Fig 1B). Although highly similar in sequence and structure, the Igs of titin perform different functions depending on their location in the titin chain. The chain consists of four main regions: the Z-disk, the I-band, the A-band and finally the M-line region. The I-band contains two signaling hubs known as the N2B and N2A region as well as an unstructured PEVK region. The A-band region consist two regions known as the D- and C-zone. The N-terminal Z-disk and C-terminal M-line Igs anchor titin in the sarcomere and support multiple protein-protein interactions. In contrast, I-band Igs are not known to interact with cellular partners, their dynamics contributing to the overall elasticity of the protein. I-band Igs can be differentially spliced creating multiple titin isoforms which regulate sarcomere compliance depending on muscle type, developmental phase or as a response to muscle damage or disease (9). Titin isoforms share constitutive I-band domains (Fig 1B, proximal and distal I-band), but distinguish themselves by their combination of differentially spliced I-band domains and specialized I-band regions (N2A and N2B). In titin, the A-band scaffolds the thick filament proteins, acting as a molecular ruler. Such a molecular giant is thought to have arisen through gene duplication events; this is supported by the occurrence of domain super-repeats in its chain (10), often in modular segments (Fig 1C). Such modular segments have been identified within the differentially spliced I-band region of titin as well as the C- and D-zones of its Aband using traditional phylogenetic tree and multiple sequence alignment analysis, in conjunction with manual interpretation (10).

**Figure 1.**
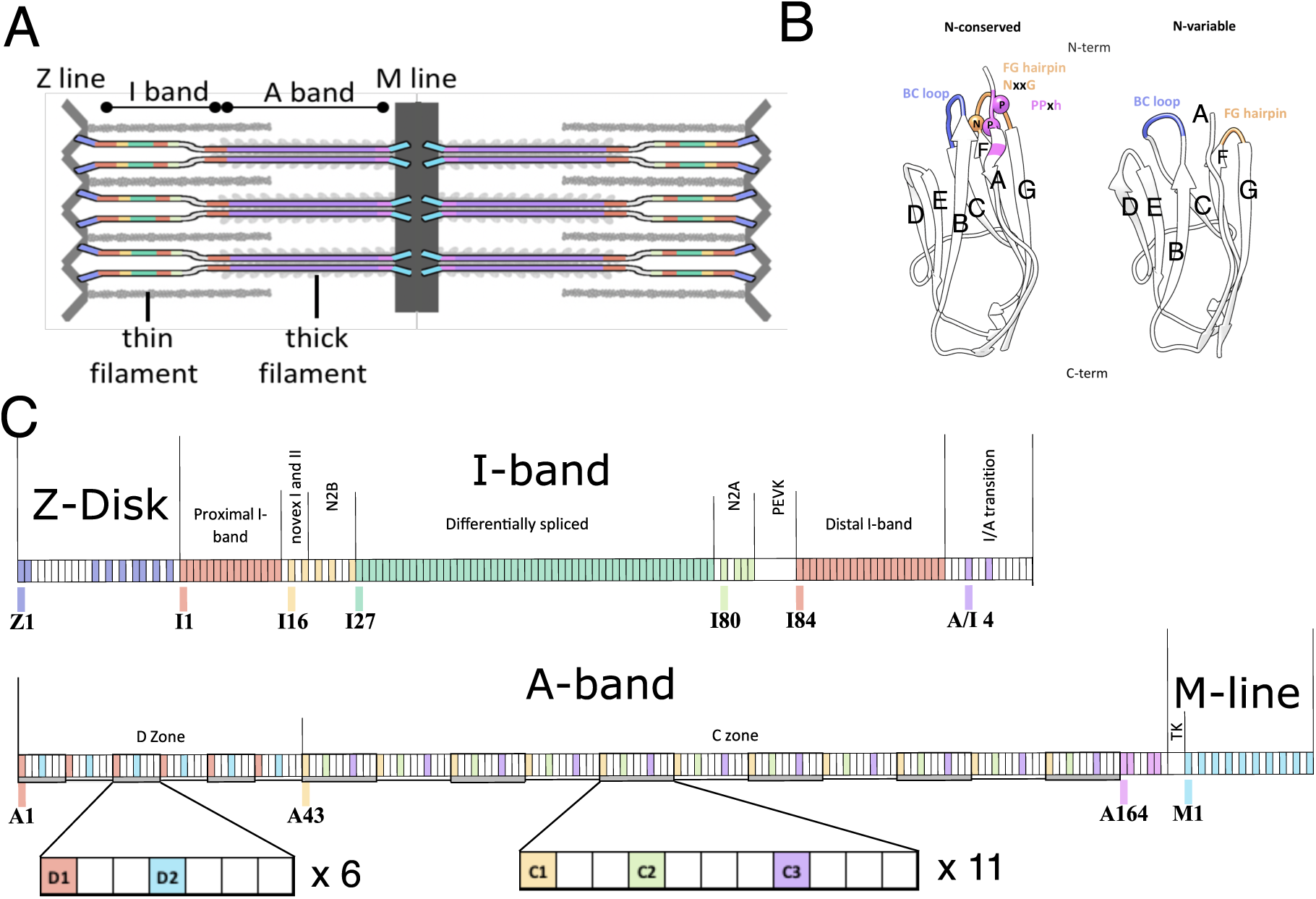
Naming convention, class features and location of Ig groups within titin. **A**. Schematic of titin showing its location in the sarcomere spanning from the Z-disk to the M-line. Titin regions are colored by Ig location: Z-line, lilac; constitutive (proximal and distal), red; differentially spliced, green; N2A, lime green; N2B, yellow; A-band, purple; M-line, cyan; pre-titin kinase (pre-TK), pink. **B**. Features of the two classes of Ig N-conserved (left) and N-variable (right) on representative structures I68 (PDB entry 2RIK) and I10 (PDB entry 4QEG). β-strand nomenclature (A-G) is shown. **C**. Location of titin Ig domains within the longest theoretical titin chain. Igs are colored as in **A**. except A-band Igs which are colored by position in each C- and D-zone repeat. Non-Ig domains are shown in white, box sizes are not representative of domain sizes. Igs at the start of each region are labeled and named according to region and domain number in this region.

Titin Ig domains have received special attention in recent years because their mutation has been linked to both skeletal muscle and heart disease (11–13) and therefore have been selected as the focus of this work. The Ig fold of titin domains is of the intermediate(I)-set type and for a number of members, their structure and sequence have been well characterized ((14) and references within). Igs in titin have traditionally been classified into subgroups according to their location in the titin chain (Fig 1A and B, Fig S1). As structural information of the domains became available, an alternative classification that accounts for sequence/structure relationships in the loop features of their Ig fold was proposed (15). In this convention, Ig domains are described as two classes, named N-conserved and N-variable, based on the features of the N-terminal pole of their fold (Fig 1C). N-conserved domains have a lengthened Ig fold characterized by a long β-hairpin between the F and G β-strands (Fig 1C, orange) containing a NxxG sequence motif, a correspondingly long proline-rich BC loop containing the motifs PPx or PxP (Fig 1C, blue) and a PPxh (h: hydrophobic residue) motif in β-strand A (Fig 1C, pink). These features pack together at the N-terminal end of the Ig fold. In contrast, N-variable Ig domains lack these features, are shorter in length and, as their name suggests, are more variable in sequence composition. Domains of the N-variable and N-conserved type correlate well with the constitutive and differentially spliced I-band regions of titin. However, this classification has proven incomplete as it fails to class several Igs that present mixed features such as domain I81 from the N2A region that has an N-conserved FG β-hairpin paired with a variable BC loop (16). The loop structure of Igs in titin is of functional significance because it affects the packing of the serially linked domains within the chain, and in this way the local chain topology, as exemplified by the 3D-structures of the N-conserved tandem I65-I70 (17), the N-variable tandem I9-I11 (18) and the mixed tandem I81-I83 (19). This in turn affects the dynamics of the chain, controlling inter-domain rotational freedom and flexibility (14), and influences sequence conservation and residue exchange tolerance at the domain-domain interfaces (20). Furthermore, an individualized loop structure in some Igs confers the domain the capability of recognizing and interacting with cellular partners. For example, this has been shown for Ig A169, where a unique loop between β-strands A and A’ is critical for the binding of the E3 ubiquitin ligase MuRF1 (21), and for I81, where an extended BC loop mediates its recruitment of the ankyrin-repeat response factor CARP (16, 22). Conversely, some Igs act as protein recruiters but do not have obvious unique features, e.g. Z1 and Z2 at the N-terminus of titin that bind telethonin (23). In these cases, it is proposed that it is the relative orientation and spacing of domains which allows for the specificity of the scaffolding (14, 23, 24). Such Ig with no specialized structural features have been often found to bind protein partners through a β-sheet augmentation mechanism that is largely sequence-independent (25). Furthermore, several additional Igs in the Z-disk (26–28), M-line (29–31), N2A (32, 33) and N2B elements (34–36) are proposed to interact with many diverse proteins. Most of these interactions currently await validation and molecular characterization. Additionally, not all unique sequence features are currently known to lead to a gain-of-function in titin Ig domains. Finally, the correct interpretation of Ig sequence features is of high importance for the interpretation of titin single nucleotide variants and their link to muscle pathology, particularly in the heart, where they are intensely studied (13).

Here, we aim to improve the classification of titin Ig domains to allow for the improved interpretation of their structural, functional and mutational features. We show that PaSiMap reassuringly separates known domains into their main classes, but is also sensitive enough to reveal subtle differences in closely related sequences, which are difficult to identify through phylogenetic trees or other clustering software. This could be generally applied in the characterization of any protein family to identify prototypical family members and those with distinguishing features and possibly divergent functions.

## MATERIAL AND METHODS

### Ig-domain protein sequences

169 titin Ig domain sequences were extracted from the translation of the inferred complete titin metatranscript (NCBI NM_001267550.2; UniProtKb Q8WZ42). This sequence contains all possible titin Igs except for I18-I23 of the short titin isoform novex 3. Domain boundaries were manually curated by alignment of all domain sequences with all titin Ig structures in the Protein Data Bank (PDB) using PROMALS3D (37). Domains were named as in (26), using their location in the sarcomere: Z for Z-disc; I for I-band; I/A for the I-to A-band transition region; A for A-band and M for M-line and their domain number within these i.e. Z1 is the first domain in the Z-line. Note that titin’s A-band also contains fibronectin domains, which are not included in this analysis. A schematic representation of all Igs and the nomenclatures used in this study is supplied in Fig S1.

### Creation of an N-feature exclusion dataset

Ig sequences were aligned using Clustal-Omega (38) and regions that define N-conserved and N-variable domain classes were removed. This was achieved by manual deletion of the FG β-hairpin motif “NxxG”, the longer BC loops and any N-terminal sequence before and including the PPxh motif (pink boxes, Fig S2). The deletions affect on average 3 to 12 residues of each sequence (4-11%).

### The PaSiMap pipeline

PaSiMap maps protein sequences as coordinates based on their pairwise similarities (Fig 2). In order to determine pairwise similarities between sequences, the tool first aligns independently each pair of sequences. The similarity of each sequence pair is represented as a number *q* in order to allow the mapping with the multidimensional scaling method *cc_analysis*.

**Figure 2.**
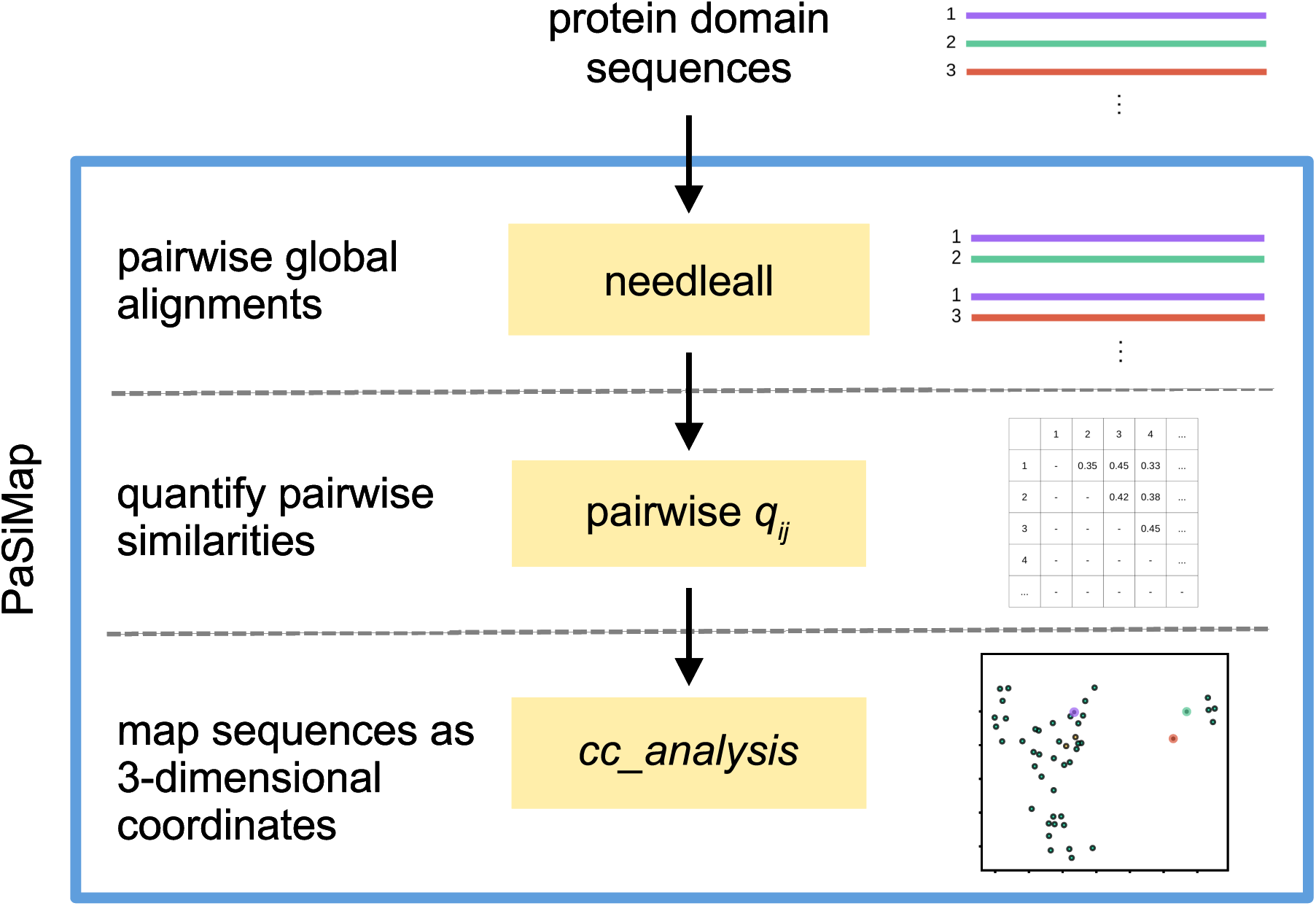
Flowchart of the PaSiMap pipeline. Blue outline: PaSiMap procedure. Yellow boxes: programs used.

### Step 1: Pairwise global alignments

Sequences were subjected to automated all-against-all global alignments using the Needleman-Wunsch implementation *needleall* of the software suite EMBOSS (39). For *N* sequences, the number of unique pairwise alignments is 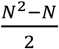 The program *needleall* was used with endweights in order to obtain pairwise alignments. All other settings were used with their default values (gapopen=10.0, gapextend=0.5, endopen=10.0, endextend=0.5, minscore=1.0, datafile=BLOSUM62).

### Step 2: Quantifying alignment similarity

Since *cc_analysis* requires numerical values for the pairwise relationships as input, it is necessary to quantify each pairwise alignment. For this purpose, we developed the quantifier *q*. This quantifier is a measure of how similar the pair of aligned sequences are in relation to the similarity that would be expected on average for a pair of randomized sequences with the same composition of amino acids.

*q* is calculated for each pairwise alignment *A*. The alignment *A* refers to the two sequences *i* and *j* : sequence *i* consists of *r*_*i*_ residues while sequence *j* consists of *r*_*j*_ residues.

Gap regions caused by insertions or deletions (i.e. where sequences are not paired) are removed from the alignment *A*. The resulting indel-free alignment *A*_*0*_ has the length *L* and consists of the aligned residues {*x*_*1*_, *x*_*2*_, …, *x*_*L*_} for sequence *i* and {*y*_*1*_, *y*_*2*_, …, *y*_*L*_} for sequence *j*.

The quantifier *q*_*0*_ is calculated for the indel-free alignment *A*_*0*_ by feature scaling:

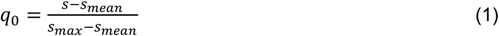

The similarity score *s* is determined with the substitution-matrix BLOSUM62 (40):

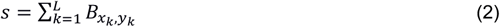

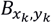: score of the substitution-matrix BLOSUM62 for the residue-pair *x*_*k*_, *y*_*k*_.

*s*_*mean*_ is defined as the similarity score which would be expected on average for a pair of randomly shuffled sequences with the same composition of amino acids as the two aligned sequences of *A*_*0*_. This calculation is deterministic and does not require explicit generation of a set of randomly shuffled sequences:

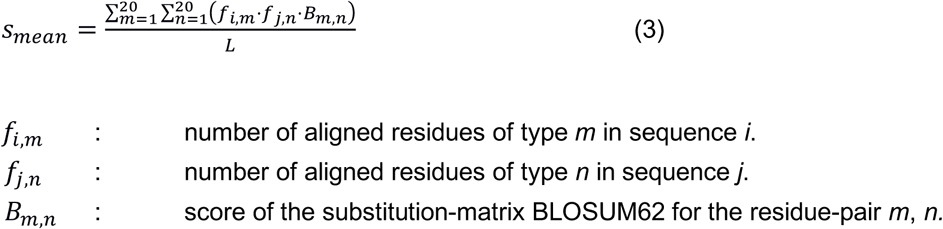

The value of *s* cannot be higher than that of a sequence aligned with itself. Thus, the maximum score *s*_*max*_ is:

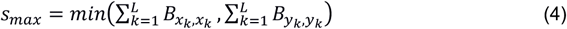

The program *needleall* only outputs pairwise alignments with score values higher than what would be expected by chance. Thus, the score *s* is always higher than the mean score *s*_*mean*_. In addition, the score *s* cannot be higher than the maximum score *s*_*max*_. Since *s-s*_*mean*_ cannot be higher than *s*_*max*_*-s*_*mean*_, the value range of the quantifier *q*_*0*_ (as calculated by equation 1) is from zero to 1.

Finally, the quantifier *q*_*0*_ is adjusted by the length of the indel-free alignment *A*_*0*_ in relation to the longer sequence of the input alignment *A*:

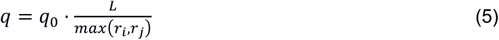

Since the coverage cannot be more than 1, the adjusted quantifier *q* has the value range from zero to 1.

In summary, short low-similarity alignments yield *q*-values closer to zero, while long high-similarity alignments result in values closer to 1. In the following, we will indicate the aligned sequences *i* and *j* by a subscript to *q*, and therefore refer to *q*_*ij*_.

### Step 3: Mapping similarity with *cc_analysis*

Previous use of *cc_analysis* (6, 41, 42) employed pairwise Pearson’s correlation coefficients as quantifiers for similarity. Pearson’s correlation coefficients have a value range from -1 to 1. However, when quantifying alignments resulting from *needleall*, negative values do not occur. Above, we defined our sequence-based quantifier *q*_*ij*_ such that its properties resemble those of a correlation coefficient in the range from zero to 1, while remaining straightforward to evaluate. The calculated pairwise relationships (*q*_*ij*_-values for each unique pair of sequences) were used as the input for *cc_analysis* (6).

Besides these pairwise *q*_*ij*_ quantifiers, *cc_analysis* also requires the user to specify the number of dimensions *d* for the output coordinates in order to iteratively, starting e.g. from random coordinates, minimize, as a function of the N vectors, the following multidimensional scaling expression:

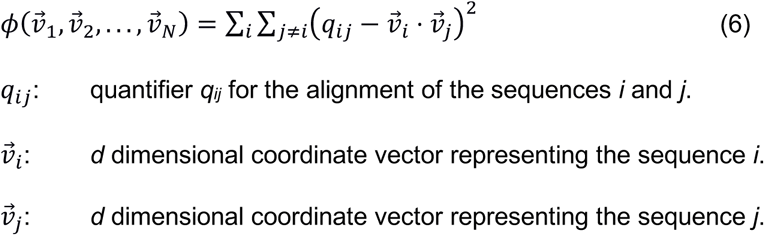

The resulting arrangement of vectors is unique except for inversion and rotation around the origin, since lengths and angles are invariant to these operations. To arrive at a standard orientation, we orient the arrangement of vectors such that the principal axes of their inertia tensor are along the axes of the *d*-dimensional coordinate system. Loosely said, this results in the first-dimension accounting for the axis with largest coordinate values, the second dimension for the second-largest coordinate values, and so on.

It was shown (6) that a) the resulting arrangement of vectors quantitatively encodes the systematic differences between the mapped objects in the angles between them, b) the length difference of colinear vectors is due to random differences of the mapped objects they represent, and c) vectors have a length of at most 1, with long vectors corresponding to a low level, and short vectors to a high level of randomness (Fig S3).

In this work, we used three dimensions (*d*=3) in order to conveniently visualize the resulting coordinates. As this limited number of dimensions might be insufficient to resolve all existent clusters, we re-analyzed (“sub-clustered”) with PaSiMap any cluster that appeared in the first pass of analysis. This is equivalent to a higher-dimensional analysis of the data, with subsequent selection of vectors that share the coordinates for their first 3 dimensions. In this work, clusters were subjected to recursive sub-analyses until no further cluster segregation was obtained.

### Plotting

*cc_analysis* represents the sequences as coordinates in low-dimensional space. These coordinates were plotted with *Matplotlib* (version 3.0.2) (43). Plotting scripts are available on GitHub (https://github.com/ksu00/pasimap).

## RESULTS AND DISCUSSION

### PaSiMap: a pipeline for grouping protein sequences by similarity

PaSiMap is a user-friendly pipeline that implements *cc_analysis* for the grouping of protein sequences by similarity. It is available online at http://pasimap.biologie.uni-konstanz.de. As input, it only requires the query sequences in FASTA format. For these, it calculates the pairwise similarities, which are then fed into *cc_analysis*. Each sequence is mapped to a coordinate in multidimensional space that can be interpreted as a vector from the origin. As in principal component analysis (PCA), dimensions are ordered by the level in which they describe features of the dataset. For easier visualization, the output dimension is commonly limited to 2D or 3D space. When the sequences are mapped as coordinates in 3D space, the first dimension (x-coordinate) represents the most influential feature of the sequences, the second dimension (y-coordinate) maps a smaller orthogonal feature, the third dimension (z-coordinate) maps an even smaller orthogonal sequence feature, and so on.

In this way, the relatedness of the sequences can be assessed through their mapping into low-dimensional space. Specifically, the angle between two vectors represents the systematic difference between the two sequences considered. Sequences whose vectors have a similar direction (i.e. the angle between them is small) commonly form a cluster. Within a cluster, the longest vectors represent the canonical, most representative sequences of that cluster. The length of a vector can be at most 1 (as shown by the unit sphere in Fig 3A). The closer to the origin a sequence is represented (*i*.*e*. the shorter its vector is), the less similar this sequence is to the ‘prototype of the group’ (*i*.*e*. longest vector with a similar angle) (Fig S3).

**Figure 3.**
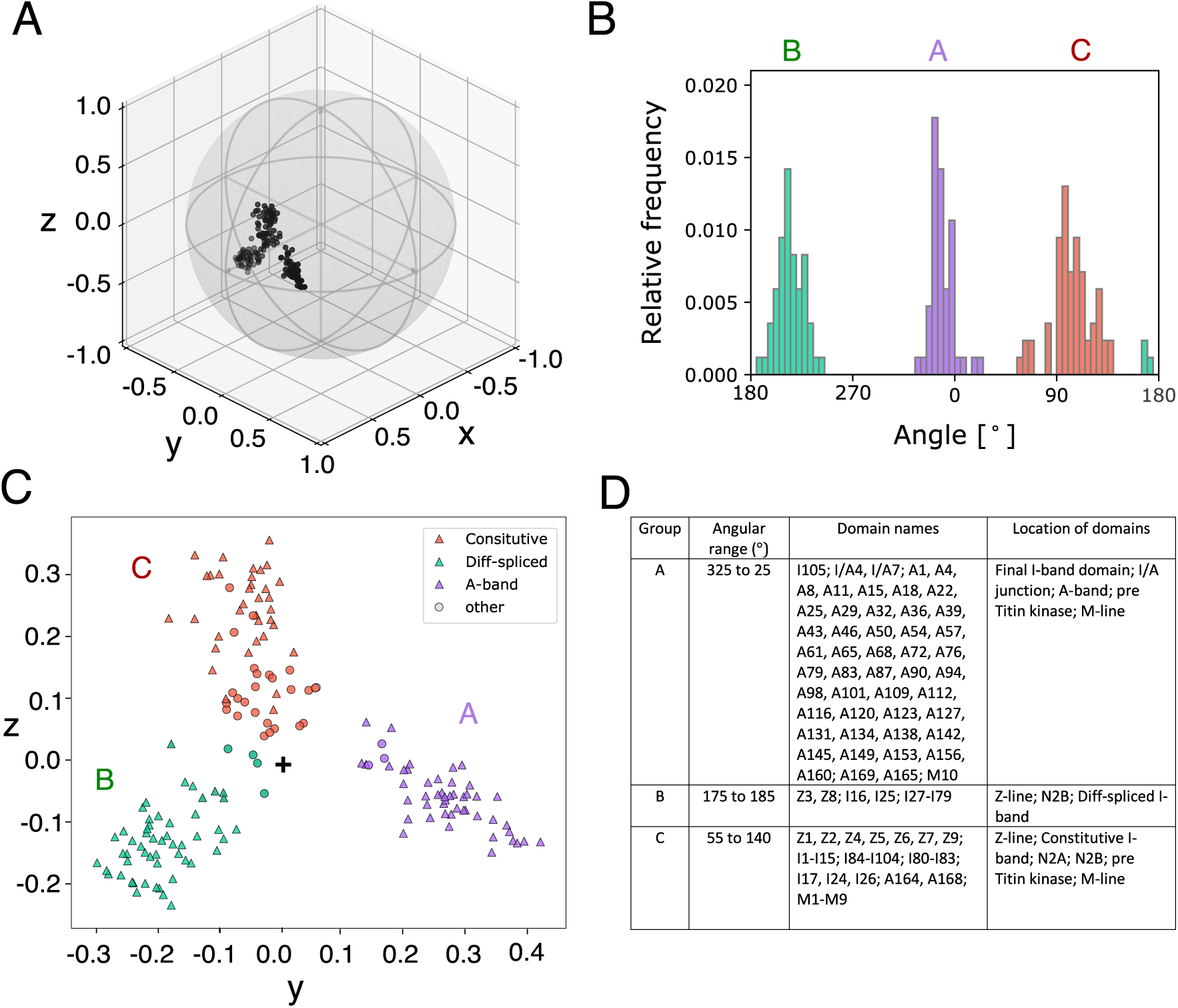
PaSiMap analysis of titin Igs reveals three major Ig groups. **A**. Mapped protein domain sequences (in 3-dimensional space). Each point represents a sequence and can be seen as a vector from the origin at (0, 0, 0). The unit sphere was added to visualize that the vectors cannot be longer than 1. **B**. Histogram for the angle distribution of the mapped protein sequences in the yz-plane. A binning size of 5° was chosen for the angles. The three biggest clusters (referenced in C) are colored in green, purple and red. **C**. Projection of the mapped sequences of **A** to the yz-plane. Ig domains of the three structural regions in titin are shown as triangles, the others as circles. **D**. Table of clusters and their protein domain members. For each cluster, the table lists the name and the location (within titin) for its members.

Without subsequent investigation, it is not possible to directly link specific sequence features to any particular dimension of the output.

### PaSiMap segregates titin Ig domains into three groups

All Ig domains in titin, amounting to 169 sequences, were subjected to the PaSiMap pipeline and mapped in 3D based on their pairwise similarities (Fig 3A). The first dimension (x-coordinate) of their vectors represents the strongest feature, namely their overall sequence similarity; these values are high but show little spread and are therefore not useful for revealing different properties of the sequences. Thus, we used a 2D plot to map the second and third dimensions which represent more subtle sequence features. The initial plot revealed three main clusters (Fig 3C). To best identify groups of sequences, the angle distribution of the sequences was visualized as a histogram (Fig 3B) that showed 3 peaks separated by more than 30°. Cluster A (purple) is mostly composed of A-band Igs, whilst Cluster B (green) is mostly composed of differentially spliced I-band domains (with a few exceptions in each case). Finally, Cluster C is composed of constitutively expressed I-band Igs and Igs from functional regions of titin such as the Z-disk, N2A and N2B segments and the M-line, many of which have attributed interactions with cellular partners (Fig S4). The separation of the constitutive I-band, differentially spliced I-band and A-band into their own groups had been proposed before using a phylogenetic analysis of Ig domains from 16 titin-like proteins (10).

It is observed that A-band Igs (purple triangles; Fig 3C) have the greatest systematic difference from the other titin Igs, even greater than the difference between spliced and constitutive Igs of the I-band. This is revealed by the differences in cluster coordinates in the second (and, therefore, hierarchically dominant) dimension, Δy, which shows that clusters B and C are vicinal while cluster A, containing A-band Ig (purple triangles), is the most distant. This indicates that A-band Igs share distinct features that distinguish them from the rest of the Igs in titin. The more pronounced unrelatedness of the A-band domains is unexpected as previously the I-band groups were distinguished from each other by the structure of their Ig folds, corresponding to the N-conserved and N-variable subtypes, respectively. Thus, the result reveals that clustering does not correlate with domain segregation according to the dominant N-pole features of the Ig fold, but that separation by location within the titin chain is more dominant, followed then by the N-pole type.

Cluster C at first inspection appears to be a heterogenous group containing Igs of the N-variable and N-conserved type, the latter appearing to be mostly Igs of the Z-disk and M-line. Because of the different fold types, it is unexpected that these Igs would form a cluster together. As discussed in Materials and Methods, the primary analysis may not display all clusters. Therefore, all clusters were further subjected to PaSiMap reanalysis to establish terminal groupings.

### PaSiMap allows for the classification of mixed and atypical feature Igs

When Cluster C is analyzed by PaSiMap it separates further into a group containing N-conserved Igs from the Z-disk, M-line and a few N-conserved types found within the constitutive I-band such as I1 (Fig 4, Cluster N-c, pink) and N-variable Igs (Fig A, Clusters Nv-1 and Nv-2, blue and yellow respectively). This further confirms the above observation that separation by location within the titin chain is the more dominant feature. The only exception is I101, which contains a NxxG loop but segregates with the other distal Igs in cluster N-v1. On the third axis (z) another cluster of N-conserved Igs can be seen (Fig 4, N-v2, green). This contains alternating Igs from the distal I-band region (I84-I104), revealing the existence of a previously undescribed repeat pattern (Fig 4C). This demonstrates the sensitivity of PaSiMap over conventional analysis procedures. I85 from this region also appears to be divergent, displaying short vector lengths in the initial PaSiMap plot, as well as the sub-cluster analysis where it resides close to the origin at the edge of group C-A.

**Figure 4.**
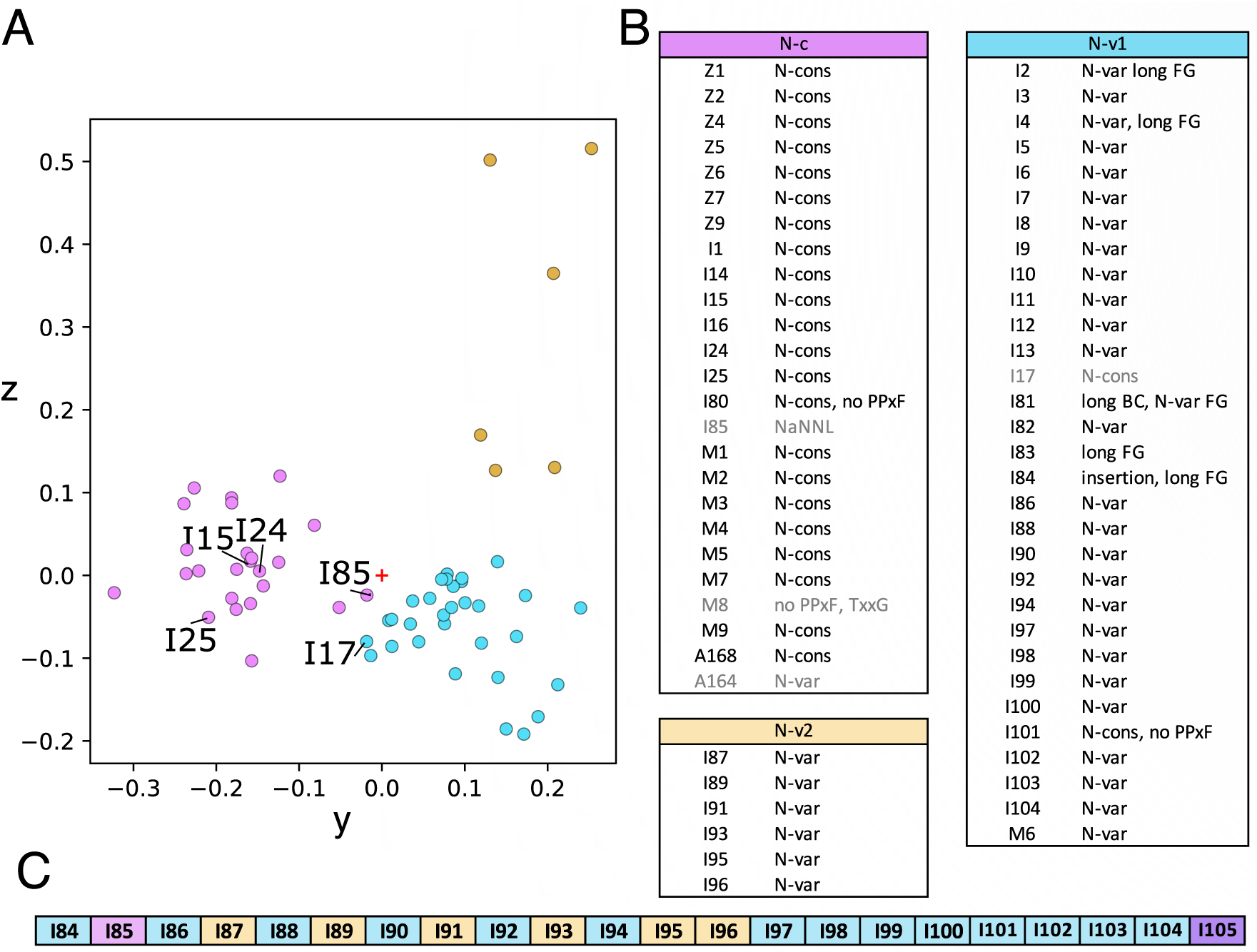
Group C subanalysis shows further separation into N-variable constitutive Igs and N-conserved functionalized Igs as well as revealing a repeat pattern in the distal I-band. **A**. Plot of domains from analysis 1 group C colored by group: N-c, pink; N-v1, blue and N-v1, yellow. Igs discussed in the text are labeled. **B**. Table of clusters and their protein domain members. Gray names indicate sequences close to the origin of the plot that could arguably be assigned to either N-c or N-v1 depending on angular cutoff used. Ig class is assigned based on the N-pole features as N-conserved (N-cons) or N-variable (N-var) with exceptions noted. **C**. Igs of the distal I-band colored by sub-analysis group, with the exception of I105 which is colored by its group in analysis 1 (Fig3).

The PaSiMap result of the N2B and N2B adjacent Igs is of particular use to analyze these domains as these contain mixed N-pole features that makes them difficult to classify. Within the N2B region Igs I15, I17 and I24 contain a NxxG motif in the FG β-hairpin and a long BC loop (even though they lack the PPxh motif in β-strand A). In the sub analysis we can see that I15 and I24 cluster with the N-conserved Igs from the M-line and Z-disk, while I17 is situated at an intermediate position (Fig4B, gray). Interestingly, domains I16 (N2B) and I25 (differentially spliced, but N2B adjacent) that also lack the PPxh motif segregate with N-conserved Igs in Cluster B in analysis 1 (Fig S4). This confirms that the features distinguishing Ig groups within titin are not restricted to the N-terminal loop structure of the Ig fold. Notable exceptions that were not detected as divergent by PaSiMap are the N2A Igs I81-I83 Igs, which are located in the middle of Cluster C in analysis 1 and remain with the N-variable Igs upon further sub-cluster analysis (analysis 2 cluster N-v1). This may be due to the small changes seen in the N-pole region and the conservation of the rest of the Ig sequence with non-functionalized Igs. This supports the hypothesis that it is the topography of the chain in non-individualized Igs which plays a role in cellular partner recognition and binding selectivity through the creation of composite binding sites involving more than one domain, such as with Z1Z2. As PaSiMap works at the primary-structure level it cannot identify such quaternary-structure level complexity.

Only a few functionalized Igs are not found in cluster C and its sub-clusters, instead locating within cluster A (hosting primarily A-band Igs). These are M10, A165, A169 and I105. As I105 is the last I-band Ig before the A band, it is unsurprising that it belongs to the A-band Ig type.

Our analysis proposes the prototypic Igs for each cluster to be A65 for cluster A (corresponding to domain 1 in super-repeat 3 of the A-band C-zone), I52 for Cluster B (differentially spliced I-band) and I88 from the heterogeneous region of titin cluster C. I88 remains as a prototypic member of Cluster N-v1, which contains mostly N-variable Igs (Fig 4, blue) and Z2 is the representative of N-c, which contains N-conserved Igs (Fig 4, pink). These Igs could be considered as prototypes in future studies of the structure and function of the diverse functional regions of titin.

### Loop differences are not the major distinguishing feature of Ig groups

To further investigate how critical the extended loop features of the N-conserved Igs are for group classification, the N-pole loops containing the features conventionally used for the classification into N-conserved and N-variable were manually removed from the input sequences (Fig S2, pink boxes, Fig S5A). The resulting core sequences were used as input in a separate analysis using PaSiMap. The clusters so calculated (Fig S5C and D, the colored underlays show the previous distribution) show that the members of the clusters remain the same. This preservation of Ig clusters confirms that the core sequence is enough to identify each Ig type and the grouping is not dominated by the sequence motifs of the N-pole loops.

### A-band C-zone super-repeats identification illustrates PaSiMap’s sensitivity

The A-band of titin is composed of well-defined domain super-repeats that are believed to underpin titin’s function as a ‘molecular ruler’, aiding the organization of the thick filament and other A-band proteins (44, 45). However, which domains are responsible for binding thick filament proteins such as myosin-binding protein C (MyBP-C) remains in contention, therefore divergent A-band Igs identified by PaSiMap will be interesting for further investigation in this area. Additionally, mutations in A-band domains are associated with various cardiomyopathies (12). The A-band consists of an A/I transition zone (which contains two Igs), six D-zone repeats of seven domains (containing two Igs here named D1 and D2), then eleven C-zone repeats of eleven domains (three of which are Igs and are termed C1, C2 and C3) and finally the pre-titin kinase domains of which four are Igs (Fig 1C). The Igs of the C-zone repeats, with a few exceptions, are known to segregate into groups depending on their position in the repeat (10). When the A-band C- and D-zone Ig domains are plotted using PaSiMap three groups that correspond to the C-zone repeats can be observed (Fig 5, yellow C1; green, C2 and purple C3). All the D-zone repeats cluster with the C3 domains (Fig 5, white circles), albeit with a wider angular spread. The C1 Igs, with the possible exception of the two first super-repeats of the C-zone A43 and A54, have been identified to bind myosin-binding protein C (MyBP-C), an important component of the thick motor filament (44, 45). However, previous analysis of their sequences in search of a difference between these domains and the other A-band Igs did not reveal any distinctive features (46). The PaSiMap analysis is clear in that the C1 Ig domains differ in a systematic manner from the C2, C3 and indeed all other A-band Igs. A43 and A54, have not been detected to be divergent by PaSiMap, and they belong clearly to the main yellow cluster with the other Igs of the C1 position. However, Tonino et al (45) suggested that it was the linker between the Ig and preceding Fn domains which lacked residues hypothesized to bind to MyBPC, and these linker residues were not included in the domain sequences given to PaSiMap. This result suggests that if there are sequence differences due to MyBP-C binding, it is less likely to be in the main body of the C1 domains, or perhaps as suggested by Bennett et al (44) this interaction is possible, but merely physically blocked by other A-band associations.

**Figure 5.**
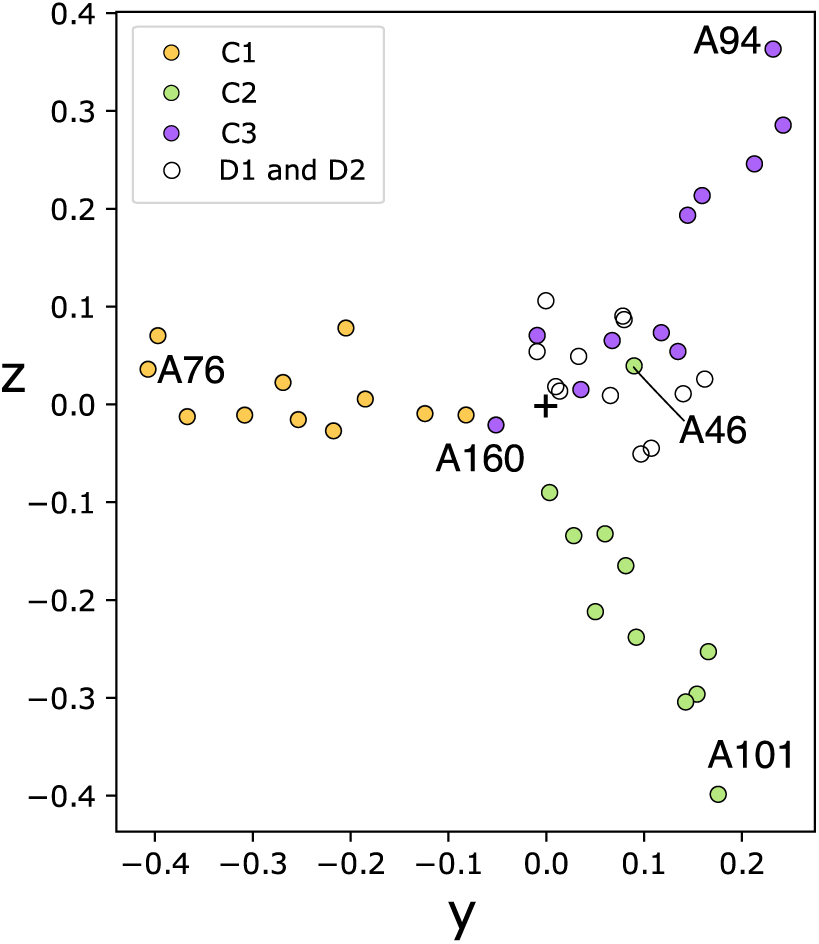
PaSiMap analysis of A-band Ig tandems. Plot of A-band domains in the C and D-zone. C-zone repeats colored as shown schematic C zone repeats in Figure 1C, D-zone Igs colored white.

There are two exceptions to this distinct clustering, A160 and A46 (Fig 5, labeled). A160 which is the final Ig of the C-zone clusters with the C1 Igs, although it has a very short vector length almost locates at the origin. This segregation of A160 with the C1 domains agrees with a previous phylogenetic analysis of titin Igs (10). A46, which is the second Ig of the C-zone, clusters with the C3 Igs. A46 was previously annotated to belong with the C2 Igs in Kenny et al (10), however, in the phylogenetic tree, it is seen to branch very close to the root of the tree and could have easily been assigned to any group. Here we see a benefit of PaSiMap allowing for an unambiguous interpretation, in comparison to phylogenetic trees where the user-defined positioning of the branches can influence the interpretation.

### Identifying repeats in titin’s differentially spliced I-band

As PaSiMap was able to identify the repeats in the A-band, we then questioned if it can also decipher the I-band repeats of the differentially spliced I-band. These repeats are less well defined. PasiMap produces four main clusters (Fig 6B yellow, blue, green and pink) and two smaller clusters (Fig 6B white and grey). As the number of mapped sequences is smaller in this case, groups were defined with a looser threshold than before (occupied angles with less than 10º separation) (Fig 6A and C; sequences close to the origin or outside of a main cluster are colored by their closest angular group but paler). When the differentially-spliced I-band Igs are colored by these groups and aligned into repeat patterns as proposed by Kenny et al 1999 (10), an overall good agreement can be seen between the two analyses. In our analysis, domains at positions 1-5 and 7-9 clearly display a repeating pattern, but domains at positions 6 and 10 show greater variability. In particular, position 6 shows the most variability, containing Igs from 5 different clusters and most with short vector lengths, indicating that they are the least similar to the representatives of each cluster and more difficult to group. This variation was not revealed in previous analysis. It is suboptimal in a repeat protein to have similar domains beside one another as, especially as folds such as the Ig fold are prone to domain ß-strand swapping and consequently misfolding. The latter is more likely should the domains have high sequence similarity and folding kinetics (47). It would therefore be interesting to investigate if the stabilities of the domains from different groups are also variable, as different folding kinetics of the respective groups would act to prevent misfolding in a serial repeat protein as titin. PaSiMap identifies that the prototypic domains to investigate from the four main groups for this type of analysis would be I69, I60, I55 and I51. Of these, the crystal structure of I69 is fortuitously already available within the I65-I70 tandem (17), which contains all Ig from all clusters except for repeat position 3 (blue). It can be concluded that the crystal structure of this tandem is highly representative of the titin chain topology at the differentially spliced I-band.

**Figure 6.**
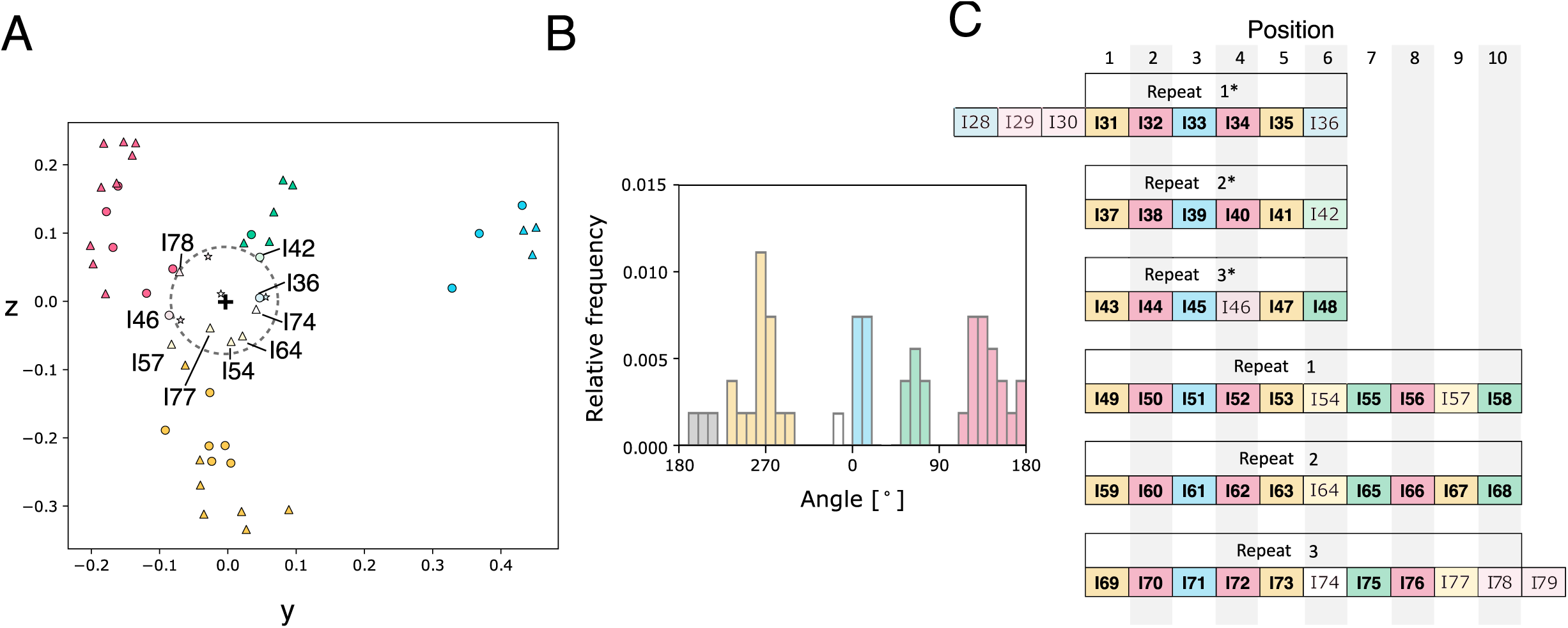
PaSiMap analysis of I-band Ig illustrates how similar sequences are rarely juxtaposed in the titin chain. **A**. PaSiMap cluster distribution of Ig components of the differentially spliced I-band tandem (I20 to I79). Sequences are colored by the groups identified in B. Sequences near the origin (grey perforated circle) are colored paler. **B**. Histogram of relative frequency of sequences by angular distribution. Bars correspond to bins of 10°. Groups have been defined as contiguously bins. **C**. Schematic of differentially spliced I-band Igs colored by PaSiMap group and repeats labeled. Igs with short vectors are colored paler.

The sub-cluster analyses of the A- and I-band illustrate that PaSiMap can highlight biologically relevant groups from a highly divergent sequence dataset such as the total titin Ig data which had as little as 6% sequence identity between some domains, as well as a sequence dataset with domains with a higher degree of similarity such as those form the differential spliced I-band and A-band Igs, which share a minimum of 15% and 17% identity and an average of 33.6% and 34.4% identity, respectively.

### New directions for titin molecular research

Structural characterization of titin Igs is highly beneficial for the interpretation of single nucleotide variants (SNVs), as it allows for *in silico* diagnostic procedures directed to clarify their disease potential (20, 48). This is a necessity for a large protein such as titin whose size renders *in vitro* and *in vivo* techniques laborious and costly. As PaSiMap excels at identifying prototypical sequences, it can be used to inform future structural and *in vitro* experimentation efforts on titin and to determine the utility of available 3D-structures from titin components for the assessment of areas of the chain with no known information. The domain group with the best structural representation to date is Cluster C, as structures are known of not only a prototypic group member, I10, but also many of the more divergent group members such as the Z- and M-line domains. The differentially spliced I-band is also well served as the tandem I65-I70 covers almost all the groups seen in the sub-cluster analysis. It is currently Cluster B containing domains from the A-band that lack appropriate structural characterization. To gain the maximum amount of insight from future structural research of this domain type and based on the current PaSiMap analysis, the following domains should be targeted: A76, A101 and A94 as these would represent the prototypic C-zone C1, C2 and C3 Ig domains (Fig 5A). The most representative A-band domain overall is A65 (Fig S6). The D-zone Igs do not have prototypic sequences but have high sequence identity to the known structures from the pre-TK region, which are represented by the structures A165 and A169 (PDB entries 2J8O and 3LCY). The final area of titin which would benefit from structural analysis is the N2B region, domains I27 and I25 in particular lack close neighbors and are likely to contribute to the specialized binding capacity of this region. Additionally, in the I-band the domain I85 has a notably short vector, despite having no known function and clustering within the region of functionalized Igs, here functional studies may be interesting to determine if it too has binding partners.

### Conclusion

Here we show that Ig domain sequence similarity in titin is hierarchical, being firstly determined by location in the protein chain, then by the N-variable or N-conserved features of their Ig fold and, finally, by the position within the corresponding super-repeat (Fig 7). The A-band analysis reassuringly reproduced previous knowledge derived about the repeats, with an improvement on the grouping of a few Igs which remained unresolved previously. The differential I-band analysis revealed that repeat position 3 is the most distinct group and position 6 represents the most diverse position in the differential I-band repeats. In addition to the established domain super-repeats in the A-band and differentially spliced I-band tandem, PasiMap reveals that a previously unidentified domain repeat exists in the distal, constitutive I-band. Igs with mixed features, which complicate their classification, have now been assigned to groups and prototypic Igs for each group identified, allowing for more directed molecular research.

**Figure 7.**
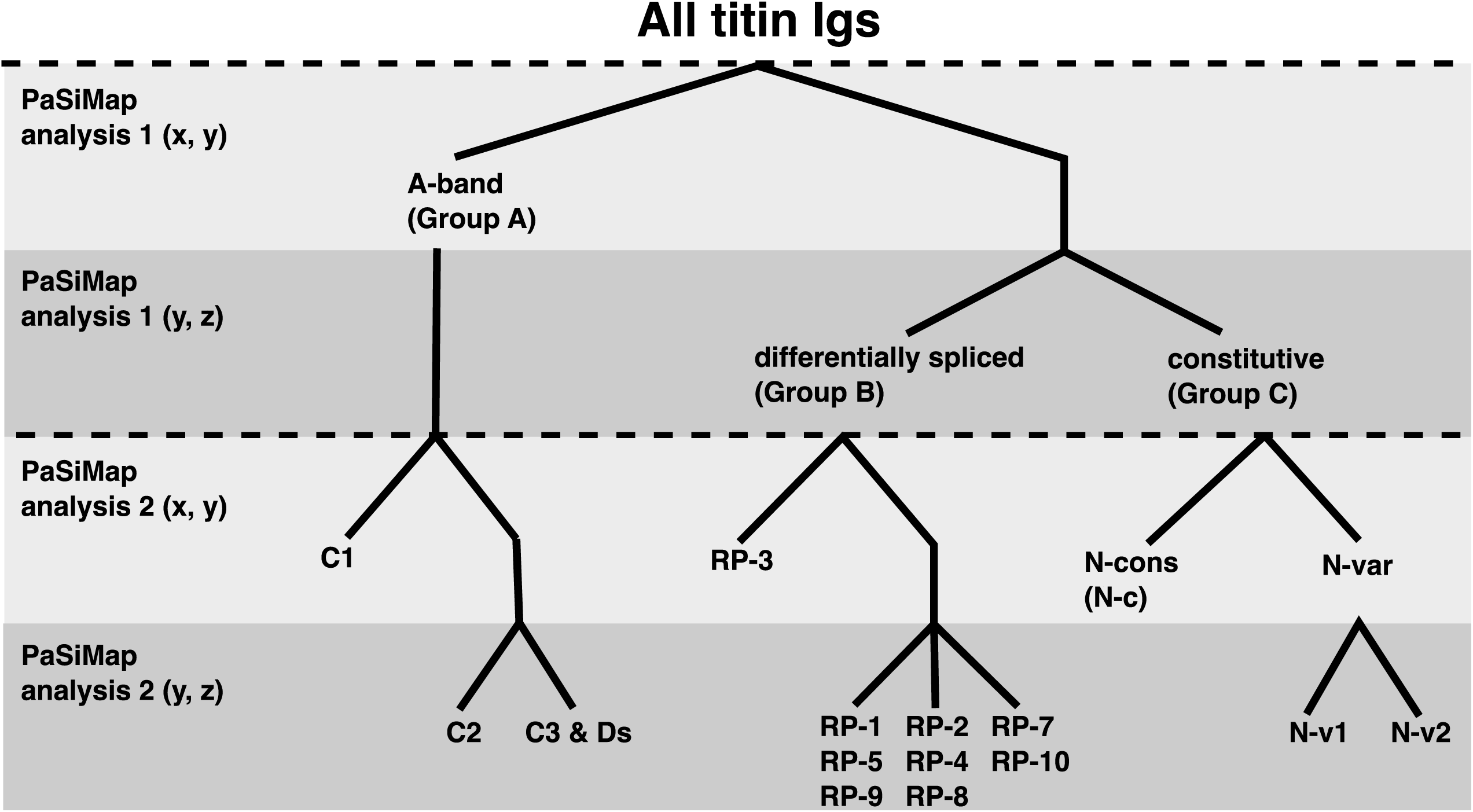
Hierarchy of cluster separation. Light gray – clusters that differentiate on the dimension 2 axis (y). Dark gray – clusters which differentiate in the third dimension (z). The first dimension (x) is not shown as the spread in this dimension is small. Top two bars correspond to PasiMap analysis 1 and bottom two correspond to Pasimap analysis 2. Clusters are named and split as follows: Group A cluster containing the A-band Igs splits into C-zone cluster C1, then C-zone 2 and 3 (C2, C3). The D-zone Igs group with the C3 Igs. Upon sub-analysis the differential spliced Igs (Group B) breaks into several clusters relating to the position of the Igs in the repeats (RP1-10), RP6 is not shown as its Igs grouped into several of the clusters (see figure 6 for details). The constitutively expressed Igs (Group C) separate into N-conserved and N-variable Igs, the latter cluster further separating into two clusters N-variable 1 and 2 (N-v1, N-v2), revealing a new repeat pattern in the distal I-band (Fig 4C).

These results show that PaSiMap is sensitive enough to distinguish small differences between similar sequences and map them in a biologically relevant way. This mapping can be used for detecting clusters of more similar sequences in a dataset, which is useful for explorative analysis. Furthermore, choosing representatives for each cluster (vector with the highest length) can be used for data reduction through the selection of a prototypic sequence for visualization or runtime-intensive downstream analyses. Another benefit of PaSiMap is that it does not directly cluster the sequences into discrete groups. Instead, the systematic differences are represented (as angles) on a continuous scale, providing additional information on which sequences are systematically more different in relation to others. This differs from phylogenetic trees which are constricted to the tree-format and linear relationships between sequences.

PaSiMap has wide applicability within sequence analysis, and is not limited to protein sequences but is also applicable to the analysis of DNA and RNA sequences. More generally, any dataset where the distinction between systematic (prototypic group representatives) and random difference (members of a unique group) would lead to insights into the biological function, would benefit from this analysis. Furthermore, the technique is not even limited to sequence analysis but can be applied to any sort of data as long as the pairwise similarity can be described as a correlation coefficient (or correlation coefficient like) value. To this end, the PaSiMap web server provides the option to input any type of pairwise relationship as correlation coefficient(-like) values instead of protein sequences. No prior knowledge of the groups is needed; just using a histogram of relative sequence frequencies, groups can be assigned in an unbiased way from the PaSiMap output. This method can therefore be used to streamline research and allow for the informed selection of representative sequences for group experiments.

## DATA AVAILABILITY

The procedure that maps protein sequences based on their pairwise similarities into low-dimensional space (as described in methods) is available as a web service PaSiMap (http://Pasimap.biologie.uni-konstanz.de/), and scripts and programs are available under a GPLv3 license (https://www.gnu.org/licenses/gpl.html) in the GitHub repository (https://github.com/ksu00/pasimap).

## SUPPLEMENTARY DATA

Supplementary Data are available at NAR online.

## ACKNOWLEDGEMENT

The authors gratefully acknowledge funding and support of KS by the Konstanz Research School Chemical Biology.

