## Supplementary Figures for "Pairwise sequence similarity mapping with PaSiMap: reclassification of immunoglobulin domains from titin as case study"

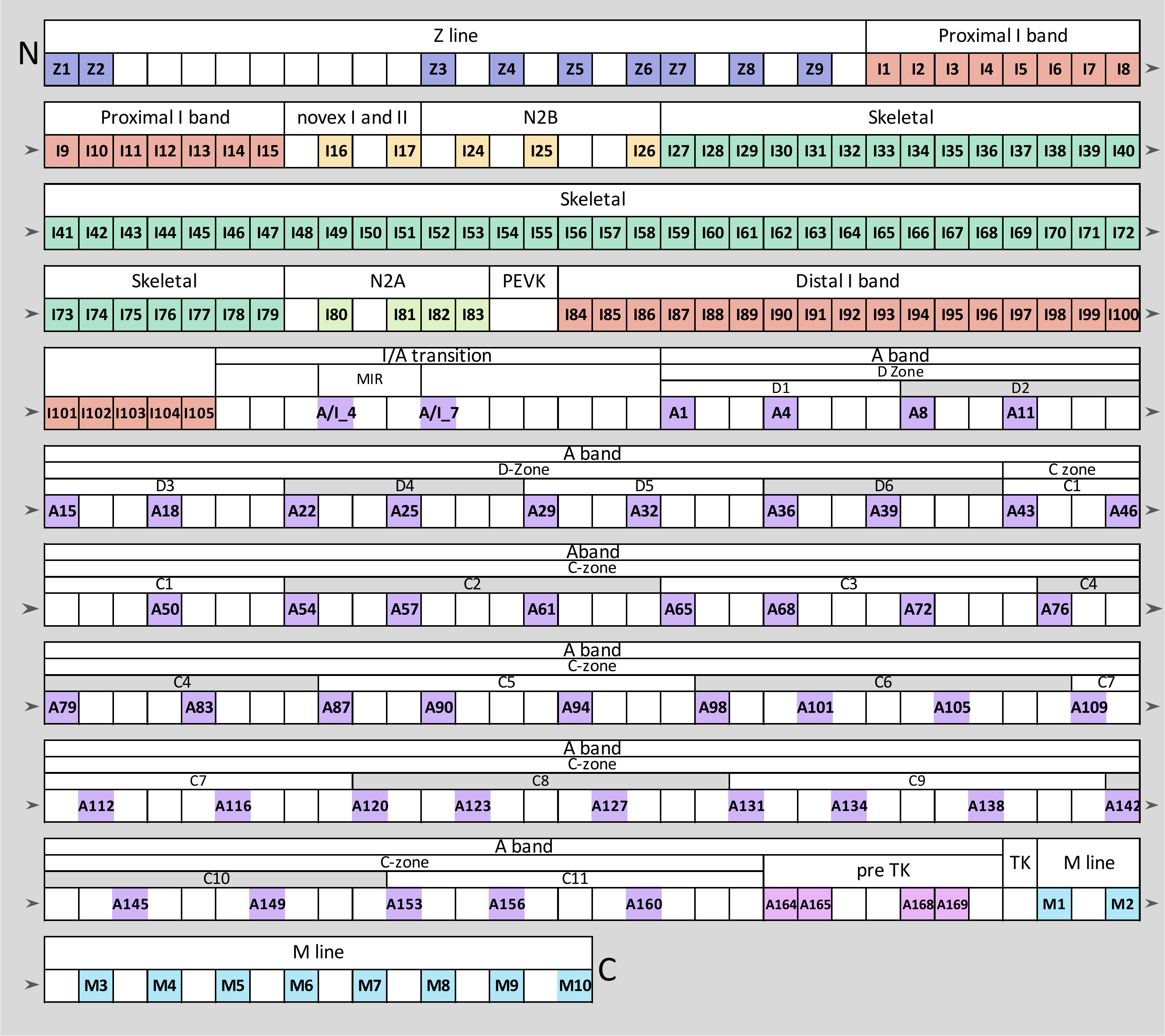


**Supplementary Figure 1. Location of titin Ig domains within the longest theoretical titin chain.**

Igs coloured by classes location: Z-line, lilac; constitutive (proximal and distal), red; alternatively spliced, green; N2A, lime green; N2B, yellow; A-band, purple; M-line, cyan; pre-Titin kinase (pre-TK), pink. Non-Ig domains are shown in white, box sizes are not representative of domain sizes. Igs are named according to region and domain number in this region.

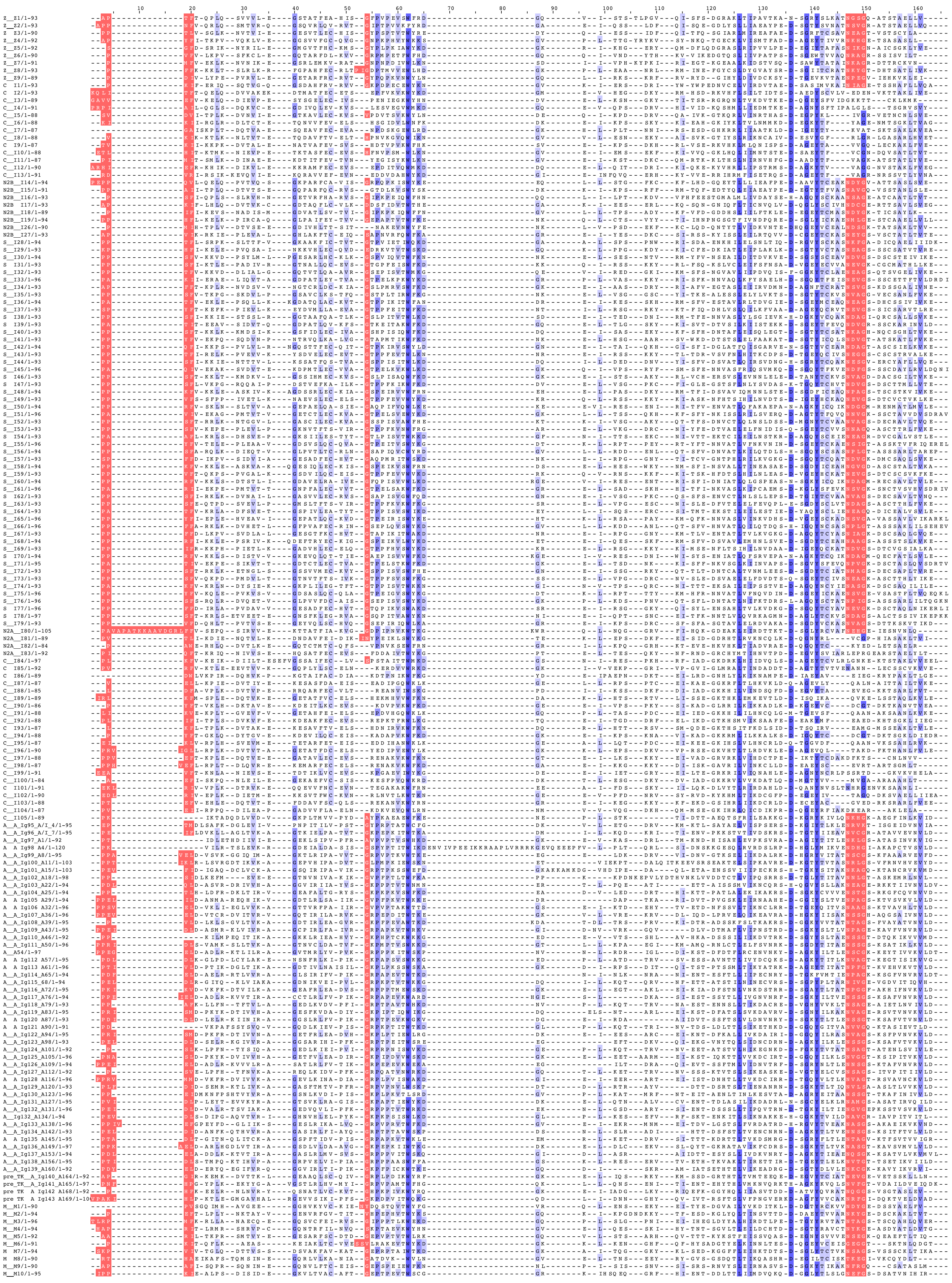

**Figure S2. Alignment of input sequences.**

N-pole loops containing the features conventionally used for the classification into N-conserved and N-variable highlighted in pink

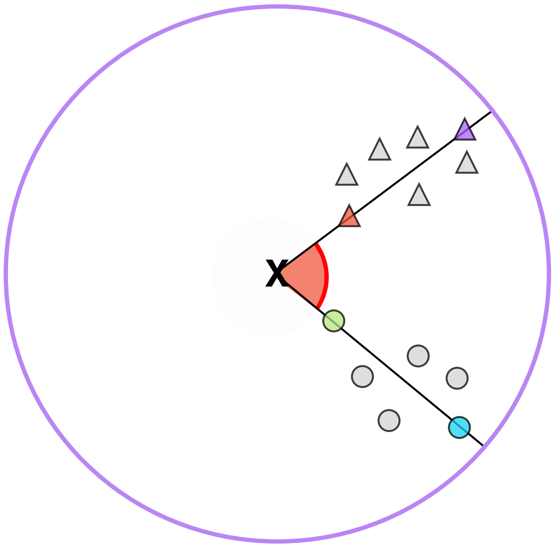


**2 groups with feature differences - circles and triangles**

- triangle group prototype
- triangle group member with the most unique differences

- circle group prototype
- circle group member with the most unique differences

**Supplementary Figure 3. Illustration of 2D PaSiMap output.**
The angle between two vectors (red semi circle) represents the systematic difference between the two sequences considered, i.e sequence features that are shared by some but not all sequences in the input group. The length difference between two vectors (with origin always fixed in the centre of coordinates (black X)) represents the unique differences between that sequence pair; these are features that are not shared by any other sequences in the analysed set. Sequences whose representing vectors have similar angles can be considered a group (here either triangles or circles). Within a group, the longest vectors represent the canonical, most representative sequences of the group (purple triangle or blue circle). The length of the vector can maximally be 1 (shown by the purple circle), The closer a sequence lies to the origin (i.e. shorter vector; red triangle or green circle) the less similar this sequence is to the ‘prototype of the group’ (i.e. longest vector with similar angle; purple triangle or blue circle).

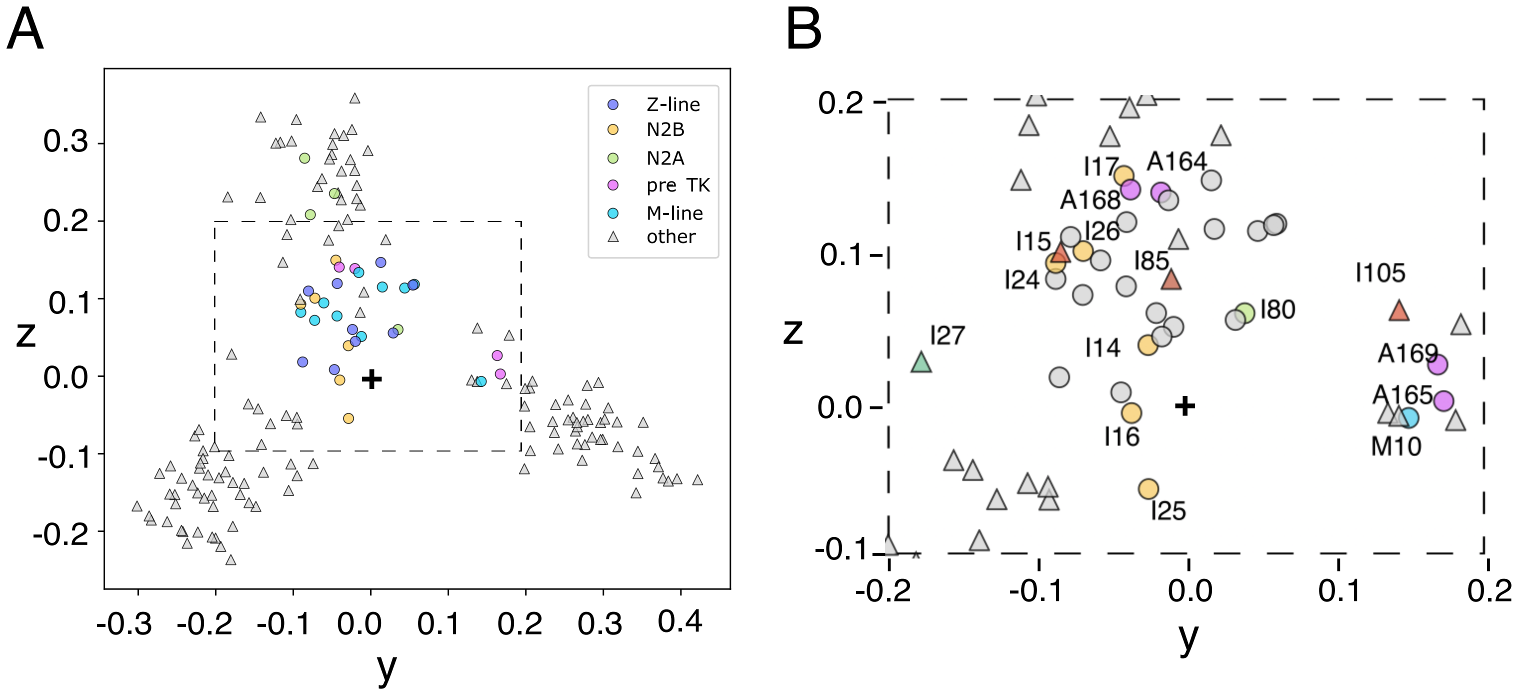


**Figure S4 Location of specilised Igs within first analysis**
**A.** First PaSiMap analysis as in Fig 3C except Igs with known or suspected specialised functions (shown as coloured circles). These are either in protein interaction regions (Z-line, slate; N2B, yellow; N2A, lime green; MuRF1 binding area, pink or M-line, cyan). **B.** Boxed area in A. zoomed in and labelled. Domains mentioned in the text are shown labelled and coloured as designated by their groups in Fig3C (for triangles) and A (for circles).

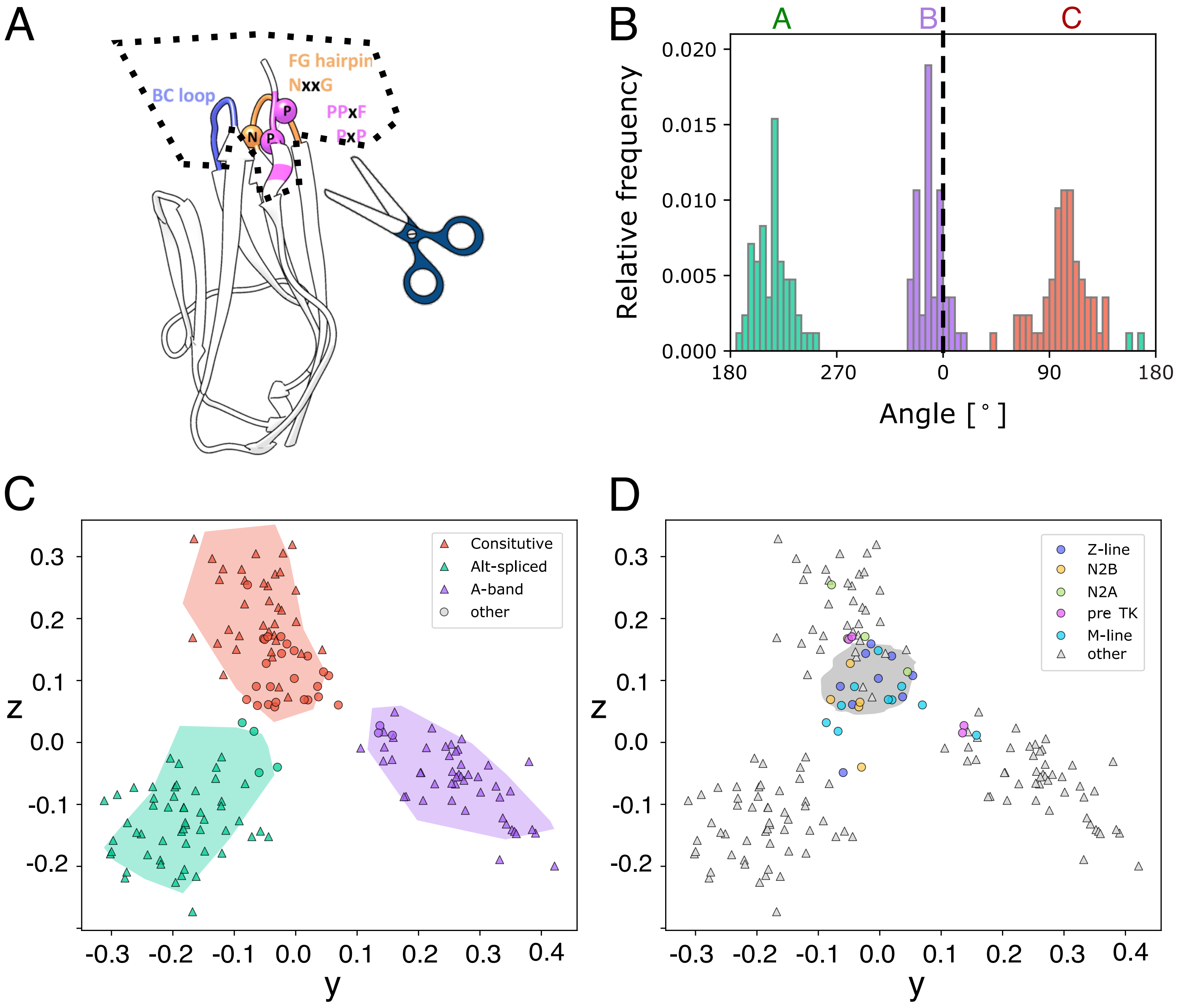


**Figure S5 PaSiMap analysis of titin Igs with N-pole features removed.**
**A.** Schematic of N-pole feature regions removed from the sequences. **B.** Histogram for the angle distribution of the mapped protein sequences in the yz-plane. The three biggest clusters (as defined in the initial analysis Fig3) are coloured in green, purple and red. **C and D.** Equivalent plots to those shown in Fig 3C and 3E with the area of the original clusters drawn underneath the new cluster distributions.

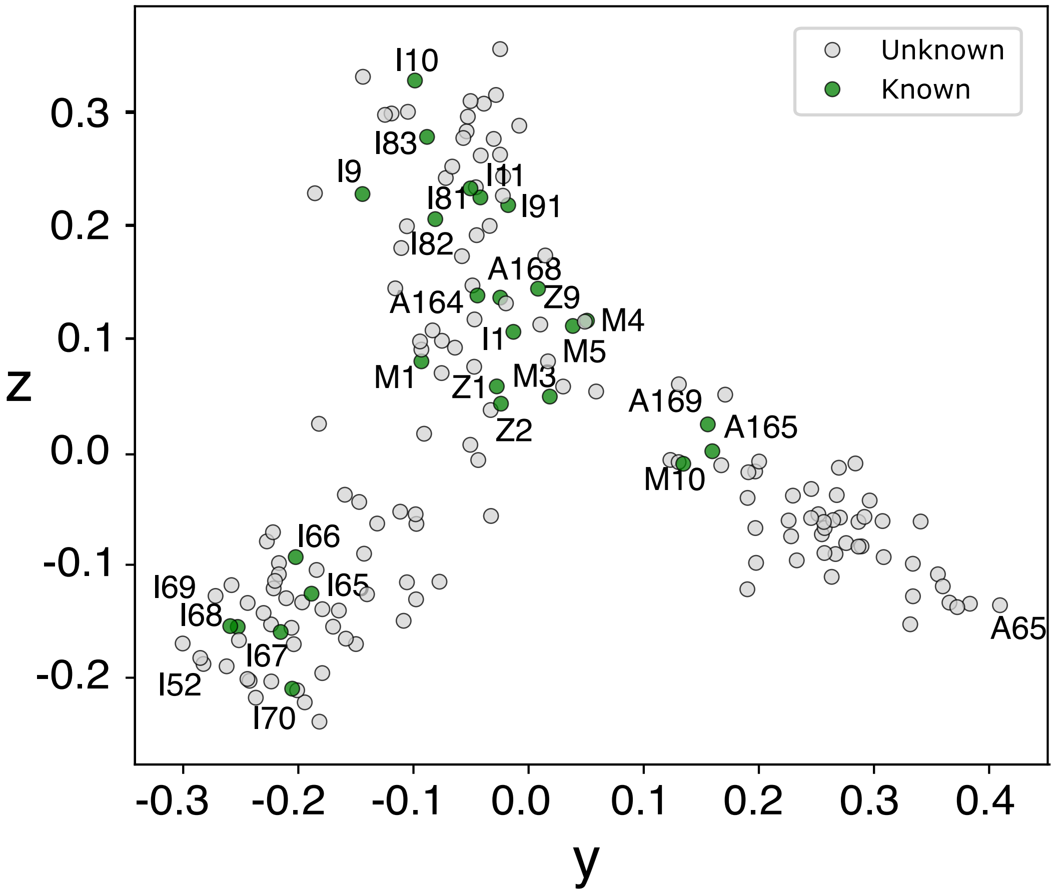


| Region | PDB code(s) |
|---|---|
| Z1-Z2 | 2A38 |
| I1 | 1G1C |
| I10 | 4QEG, 5JDJ |
| I9-11 | 5JDE, 5JDD |
| I67-I69 | 2RIK |
| I65-70 | 3B43 |
| I81 | 5JOE |
| I81-I83 | 7AHS |
| I83 | 6YJ0 |
| I94 | 1TIU, 1TIT, 2RQ8, 1WAA |
| A164-165 | 3LCY |
| A168-1169 | 2J8H, 2ILL |
| M1 | 2BK8 |
| M3 | 6HCI |
| M4 | 3QP3 |
| M5 | 1NCT,1NCU, 1TNM, 1TTN |
| M7 | 3PUC |
| M10 | 2Y9R,3Q5O |

**Figure S6. Location of domains with known structures shown in green.**
